# Post-translational cleavage of Mei5 of Dmc1 mediator, Mei5-Sae3 complex, in yeast meiosis

**DOI:** 10.1101/2024.04.25.591189

**Authors:** Stephen Mwaniki, Priyanka Sawant, Osaretin P. Osemwenkhae, Yurika Fujita, Masaru Ito, Asako Furukohri, Akira Shinohara

**Author notes:** Corresponding author: Akira Shinohara Institute for Protein Research, The University of Osaka 3-2 Yamadaoka, Suita, Osaka 565-0871 JAPAN. Contributed equally (Stephen Mwaniki, Priyanka Sawant). University of Benin, Nigeria (Osaretin P. Osemwenkhae).

## Abstract

Interhomolog recombination in meiosis is mediated by the Dmc1 recombinase. The Mei5-Sae3 complex of *S. cerevisiae* promotes Dmc1 assembly and functions with Dmc1 for homology-mediated repair of meiotic DNA double-strand breaks. How Mei5-Sae3 facilitates Dmc1 assembly remains poorly understood. In this study, we created and characterized several *mei5* mutants featuring the amino acid substitutions of basic residues. We found that Arg97 of Mei5, conserved in its ortholog, SFR1(complex with SWI5), RAD51 mediator, in humans and other organisms, is critical for complex formation with Sae3 for Dmc1 assembly. Moreover, the substitution of Arg117 with Ala in Mei5 resulted in the production of a C-terminal truncated Mei5 protein during yeast meiosis. Notably, the shorter Mei5-R117A protein was observed in meiotic cells but not in mitotic cells when expressed, suggesting a unique regulation of Dmc1-mediated recombination by post-translational processing of Mei5-Sae3.

## Introduction

Meiotic recombination plays a critical role in chromosome segregation in meiosis I and in generating a new combination of alleles in gametes for diversity (Marston, 2014). The recombination promotes reciprocal exchange between homologous chromosomes creating physical linkages between them, rather than between sister chromatids (Cejka, Mojas, Gillet, Schar, & Jiricny, 2005; Hunter, 2015). This interhomolog bias is regulated by modulating the recombination machinery as well as chromosome structure during meiosis.

Meiotic recombination in the budding yeast, *S. cerevisiae*, is induced by the formation of DNA double-strand breaks (DSBs), followed by the generation of single-stranded DNA (ssDNA) through end processing. Two RecA homologs, Rad51 and Dmc1 (Bishop, Park, Xu, & Kleckner, 1992; A. Shinohara, Ogawa, & Ogawa, 1992) form a nucleoprotein filament on the ssDNA for homology search and strand exchange between the ssDNA and a homologous double-stranded DNA (dsDNA) (Luo et al., 2021; Ogawa, Yu, Shinohara, & Egelman, 1993; Xu et al., 2021). In meiosis, Rad51 plays a non-catalytic, structural role in interhomolog recombination by aiding the assembly of Dmc1 filament (Cloud, Chan, Grubb, Budke, & Bishop, 2012; Lan et al., 2020) while Rad51 catalyzes the intersister strand exchange in mitosis. The Dmc1 filament catalyzes the interhomolog strand exchange. The assembly of Rad51 filament on ssDNAs coated with a ssDNA-binding protein, Replication protein A (RPA) is tightly regulated by Rad51 mediators: Rad52, Rad55-Rad57, and Psy3-Csm2-Shu1-Shu2 in the budding yeast (Sasanuma et al., 2013; A. Shinohara & Ogawa, 1998; Sung, 1997). On the other hand, Dmc1 assembly on RPA-coated ssDNAs is promoted by the meiosis-specific complex, Mei5-Sae3, as well as Rad51 (Chan, Zhang, Weissman, & Bishop, 2019; Ferrari, Grubb, & Bishop, 2009; Hayase et al., 2004; Tsubouchi & Roeder, 2004).

The budding yeast Mei5-Sae3 belongs to a family of protein complexes along with Sfr1-Swi5 in the fission yeast, which regulates the assembly of both Rad51 and Dmc1 filaments (Akamatsu, Dziadkowiec, Ikeguchi, Shinagawa, & Iwasaki, 2003). Sfr1-Swi5 and Mei5-Sae3 complexes are conserved from yeast to mammals (Akamatsu et al., 2007; Hayase et al., 2004; Tsai et al., 2012). Fission yeast and mouse Sfr1-Swi5 promote both Rad51-mediated strand exchange *in vitro* by stabilizing an active ATP-bound form of Rad51 filaments (Chi et al., 2009; Haruta et al., 2008). Consistent with this, structure studies showed that the C-terminal region of Sfr1 (Sfr1C), when complexed with Swi5, forms a kinked rod, that may fit into a groove of the Rad51 filament (Kokabu et al., 2011; Kuwabara et al., 2012). Swi5 and Sae3 encoding small proteins with fewer than 100 amino acid (aa), are homologous to each other (Akamatsu et al., 2007; Hayase et al., 2004). On the other hand, Sfr1/Mei5 is highly divergent showing limited homology (Hayase et al., 2004). Still how the formation of the Mei5-Sae3 complex is regulated in a meiotic cell is largely unknown.

Given that Mei5-Sae3 binds to ssDNA (Ferrari et al., 2009; Say et al., 2011), we constructed several *mei5* mutants in which a positively-charged amino acid is replaced with neutral amino acids, alanine or leucine, and characterized the mutants. We found that the substitution of Arg117 with alanine produced a C-terminal truncated Mei5 protein (Mei5-R117A), suggesting the presence of post-translational processing of the mutant Mei5 protein. The role of the post-translational processing is discussed.

## Results

### Mutational analysis of Mei5

Previous study showed a crystal structure of fission yeast Swi5 complex with Sfr1C (N-terminal truncated version of Sfr1) (Kuwabara et al., 2012) (PDB, 3viq). Amino acid sequence comparison revealed weak similarity between the budding yeast Mei5 and fission yeast Sfr1 (Hayase et al., 2004). In the sequence, regions of α1 and α2 of the Sfr1 protein are more homologous to those in Mei5. α1 and α2 of the Sfr1 protein form parallel α-helixes with two α-helixes of Swi5 (Kuwabara et al., 2012). Indeed, the AlphaFold2 prediction showed that Mei5 contains four α-helixes followed by two short α-helixes (Figure 1A). Two α-helixes (α1 and α2) of the N-terminal of Mei5 share the structural similarity to the two helixes of Sfr1 (Figure 1B). On the other hand, the predicted C-terminal region of Mei5, which contains anti-parallel α-helixes followed by two short α-helixes is different from the C-terminal Sfr1 with anti-parallel β-sheet followed by a short α-helix (Figure 1A).

**Figure 1.**
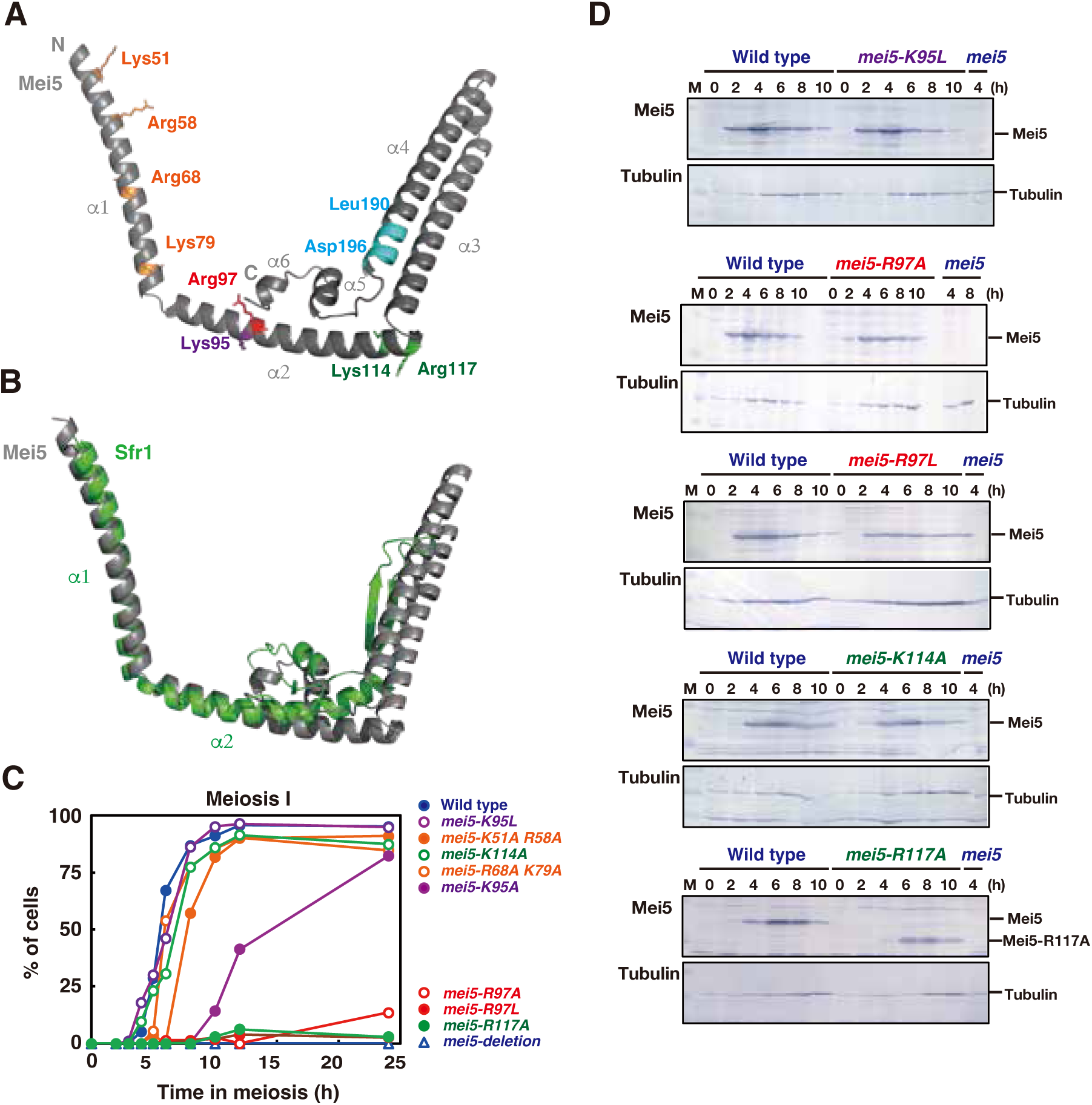
Predicted Mei5 structure and the relationship with Swi5-Sfr1. A. Predicted structure of the budding yeast Mei5 (gray) by the AlphaFold2 (https://alphafold.ebi.ac.uk); AF-P32489. The amino acid residues studied in this study are shown as a stick with a color. B. Structural comparison of a predicted Mei5 (gray) with Sfr1(green) complex. The structure alignment was performed by PyMOL. C. The entry into meiosis I in various strains was analyzed by DAPI staining. The number of DAPI bodies in a cell was counted. A cell with 2, 3, and 4 DAPI bodies was defined as a cell that passed through meiosis I. The graph shows the percentages of cells that completed MI or MII at the indicated time points. Strains used are as follows: Wild-type, MSY832/833; *mei5*::*URA3*, PSY64/65; *mei5-K51A R58A*, PSY33/34; *mei5-R68A K79A*, PSY23/24; *mei5-K95A*, PSY11/12; *mei5-K95L*, PSY123/124; *mei5-R97A*, PSY15/16; *mei5-R97L*, PSY13/14; *mei5-K114A*, PSY33/34; *mei5-R117A*, PSY78/79. More than 100 cells were counted at each time point. D. Expression of various mutant Mei5 proteins in meiosis. Lysates obtained from the cells at various time points during meiosis were analyzed by western blotting using anti-Mei5 (upper) or anti-tubulin (lower) antibodies.

We were interested in the function of the conserved α-helixes, the α1 and α2 of Mei5. Particularly, the α2 bears highly conserved KWR/K motif among Sfr1/Mei5 orthologues. We confirmed a purified Mei5-Sae3 binds to ssDNAs (Ferrari et al., 2009; Say et al., 2011). We focused on positively-charged amino acids in the α1 and α2 of Mei5; Lys51, Arg58, Arg68, Lys79, Arg90, Lys91, Lys95, Arg97, Lys114, and Arg117 (Mei5 in SK1 strain is one amino acid shorter than that in other strains-Glu17, Glu18 in SK1 instead of Glu17, Glu18, Glu19 in others) and Lys95 and Arg97 are in the KWR/K motif (Figure 1A, left). We substituted the lysine or arginine with either alanine or leucine in 7 (9) mutants; *mei5-K51A R58A* (double mutant), *mei5-R68A K79A* (double mutant), *mei5-R90A K91A* (double mutant), *mei5-K95A* (and *-K95L*), *mei5-R97A* (and *-R97L*) *mei5-K114A*, and *mei5-R117A* in the SK1 background. The staining of the cells with a DNA-binding dye, DAPI, reveals that, after synchronous induction of meiosis, *mei5-R68A K79A*, *mei5-R90A K91A*, *mei5-K95L*, and *mei5-K114A* showed the similar timing of the entry of meiosis I to that in the wild-type cells (Figure 1C). The *mei5-K51A R58A* showed ∼1 h delay in the entry into MI. The *mei5-K95A* mutant cells delayed ∼4 h in the entry in meiosis I while the *mei5-K95L* cells showed normal MI progression. Moreover, except for the *mei5-K95A,* the mutant cells expressed Mei5 proteins with expression kinetics similar to wild-type Mei5 protein, which appears at 2 h and increases in its level in further incubation (Figure 1D and Supplementary Figure S1C). The *mei5-K95A* delayed the expression of Mei5 with a reduced level relative to the wild-type (Figure 1D). The *mei5-K95L* mutant followed a similar expression pattern to the wild-type Mei5. The delayed meiosis I in the *mei5-K95A* would be due to the delayed expression of the Mei5-K95A protein (Figure 1D). Although the *mei5-K51A R58A* showed a slight delay in MI progression, the spore viability of the *mei5-K51A R58A* mutant is at a wild-type level with 97.0% (97.8% in wild-type). The spore viability of the *mei5-K95A* mutant is 96.5%.

We further characterized *mei5-K95L*, *-R97A*, *-R97L*, and *-R117A* mutants in more detail.

#### The *mei5-K95L* mutant

To know the role of highly-conserved Lys95 among Mei5/Sfr1 (see above), we further characterized the *mei5-K95L* mutant, which shows normal expression of Mei5 protein in detail. Moreover, immuno-staining analysis of Rad51 on chromosome spreads revealed the normal appearance of Rad51-positive spreads (Figure 2A) but a slight delay in the disappearance of Rad51. The *mei5-K95L* mutant delayed the appearance and disappearance of Dmc1 slightly compared to the wild-type. The peak value of Dmc1-positive cells at 4 h is lower than that in the wild-type (Figure 2C). The Dmc1-focus number in the mutant is lower than that in the wild-type (19.1 in the mutant at 4 h while 33.0 in the control; Figure 2D and Supplementary Figure S2B). These indicate that the *mei5-K95L* mutant bears a weak defect in the Dmc1 assembly. Mei5 staining revealed that the Mei5 foci appeared with a slight delay, peaked at 4 h, and disappeared at the normal timing. The number of Mei5 mutant foci is 21.4 at 4h, which is lower than that of wild-type Mei5 foci (29.4). Given that the *mei5-K95L* mutant slightly decreased the spore viability to 91.7% compared to the control (97.8%), K95 is not essential for Mei5 function *in vivo*. A non-essential role of the highly-conserved residue(s) is also seen in Y56 and N57 of the conserved YNEL sequence of the Mei5 partner, Sae3 (Sawant et al., 2023).

**Figure 2.**
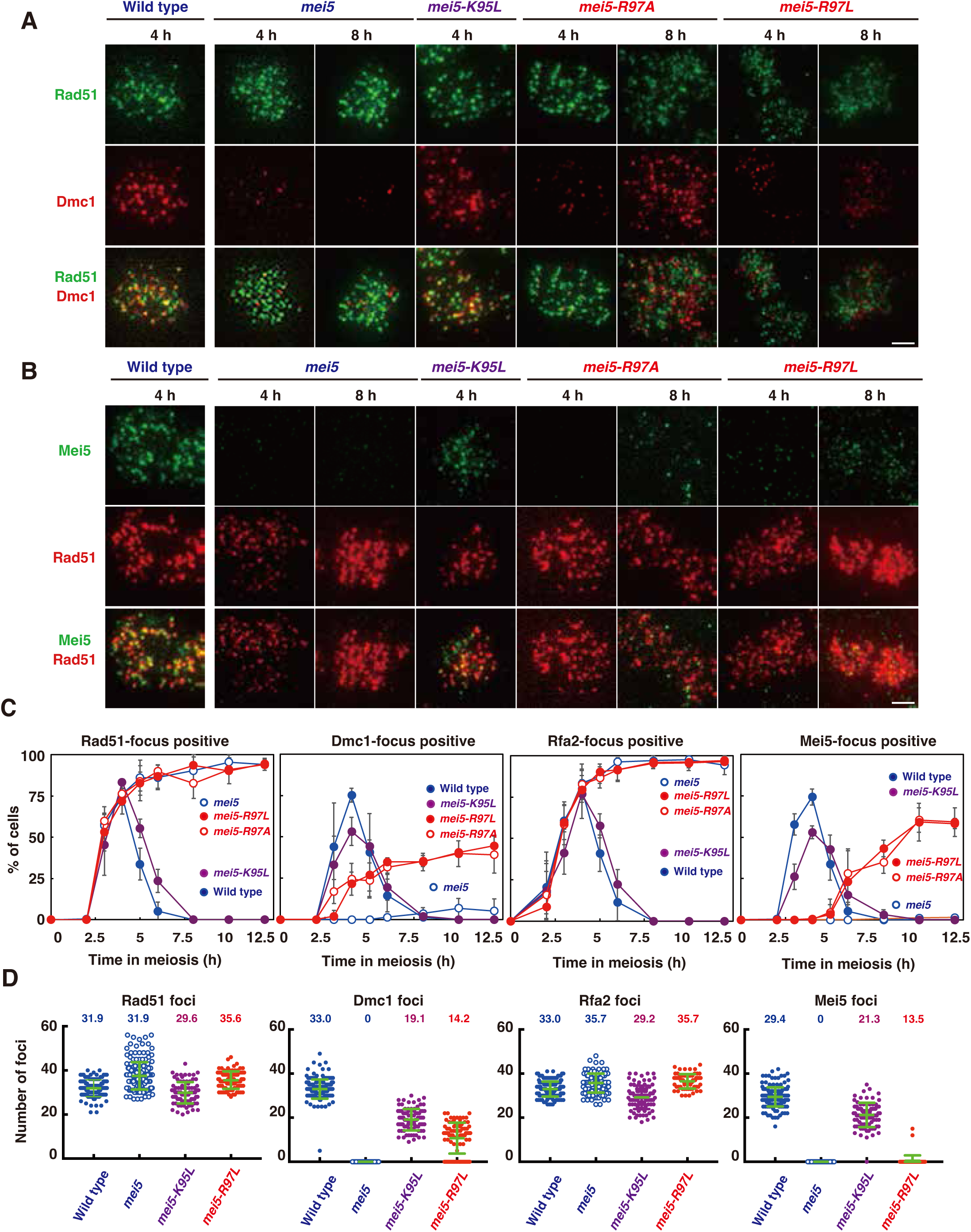
*mei5* mutations cause defective in Dmc1 assembly. A. Rad51 and Dmc1 staining. Nuclear spreads were stained with anti-Rad51 (green), anti-Dmc1 (red), and DAPI (blue). Representative images at each time point under the two conditions are shown. Strains used are as follows: Wild-type, NKY1551; *mei5::URA3* deletion, PSY5/6; *mei5-K95L*, PSY133/137; *mei5-R97A*, PSY86/90; *mei5-R97L*, PSY131/141. Bar = 2 μm. B. Rad51 and Mei5 staining. Nuclear spreads were stained with anti-Rad51 (red), anti-Mei5 (green), and DAPI (blue). Representative images at each time point under the two conditions are shown. Bar = 2 μm. C. Kinetics of assembly/disassembly of Rad51, Dmc1, Rfa2 and Mei5. The number of cells positive for foci (with more than 5 foci) was counted at each time point. At each time point, more than 100 cells were counted. The average values and SDs of triplicates are shown. D. The number of foci of Rad51, Dmc1, Rfa2, and Mei5 at 4 h was manually counted. The graphs show the focus number combined from three independent time courses. On the top, an average focus number in positive nucleus is shown. Error bars (green) is a mean with standard deviation.

### The *mei5-R97L* mutant were partially deficient in Dmc1 assembly

The *mei5-R97A* and *-R97L* mutants showed an arrest in meiotic prophase I (Figure 1C). Like the *mei5* deletion mutant, these mutants formed Rad51 foci normally like in the wild-type strain but accumulated the foci as meiosis progresses, suggesting stalled recombination (Figure 2). This is confirmed by the accumulation of Rfa2 (the second subunit of RPA) foci (Supplementary Figure S2A). The kinetics of Rfa2 foci in both of the mutants are similar to those of Rad51 foci. Moreover, the chromosomal fragmentation analysis by the CHEF (Counter-clamped homogeneous electric field) gel electrophoresis confirmed the accumulation of fragmented chromosomes induced by meiotic DSBs in the *mei5-R97L* mutant as seen in the *mei5* deletion mutant (Supplementary Figure S3C).

Dmc1 staining revealed that the *mei5-R97A* and *-R97L* mutants are partially defective in Dmc1-focus formation (Figure 2C and D). At early time points such as 3 and 4 h, the appearance of Dmc1 foci in both mutants is delayed relative to the wild-type control. Moreover, at 10 and 12 h, only ∼40% of mutant cells were positive for the Dmc1 foci, suggesting a defect in Dmc1 disassembly. The steady-state number of Dmc1 foci per spread in the *mei5-R97L* mutant is 10.7 and 14.4 at 4 and 8 h, respectively while that in the wild-type is 33.0 at 4 h (Supplementary Figure S2B), consistent with that the *mei5-R97L* mutant is defective in Dmc1 assembly. The mutant accumulated Dmc1 foci at late times in meiosis. Therefore, the Dmc1 complex formed in the *mei5-R97A* and *-R97L* cells is likely to be functionally defective.

The Mei5 staining revealed a defective assembly of Mei5 in the *mei5-R97A* and *-R97L* cells (Figure 2B, C and D). At early time points such as 3, 4, and 5 h, we could not detect a clear signal of Mei5 on the spreads. After 6 h, Mei5 foci started to appear and accumulated with ∼50% of cells positive for Mei5 during further incubation. On the other hand, the number of Mei5 foci in the *mei5-R97L* mutant was 10.2 and 9.8 at 6 and 8 h, respectively, which is lower than that of Mei5 foci in the wild type (29.4 at 4 h). These suggest that the Dmc1 assembly in the *mei5-R97A and -R97L* mutants occurs without visible Mei5-focus formation. Moreover, it seems that the mutant Mei5-R97L/A protein could bind to a pre-assembled Dmc1 ensemble.

### Arg97 in Mei5 protein is required for efficient complex formation with Sae3

To know the interaction of the Mei5-R97L protein with Sae3, we introduced the *SAE3-Flag* allele, which is functional (Hayase et al., 2004; Sawant et al., 2023), in the *mei5-K95L* and *mei5-R97L* mutants. Like the *mei5-R97L*, *mei5-R97L SAE3-FLAG* cells, which expressed both Mei5 and Sae3 protein as seen in the wild-type (Supplementary Figure S3A) were arrested at meiosis I and accumulated Rad51 and Rfa2 foci (Figure 3C, Supplementary Figure S3B). As seen in the *mei5-R97L*, *mei5-R97L SAE3-FLAG* mutant accumulated meiotic DSBs (Supplementary Figure S3D), indicating defective processing of recombination intermediates. Distinct from *mei5-R97L*, the *mei5-R97L SAE3-FLAG* mutant is almost defective in Dmc1-focus formation (Figure 3A and C; compare with Figure 2A), indicating a genetic interaction between the *mei5-* and *SAE3-Flag* alleles. The number of Dmc1 foci in the mutant is 2.3 and 3.0 at 6 and 8 h (Figure 3D). Moreover, the focus formation of both Mei5 and Sae3-Flag is fully impaired in the mutant (Figure 3B).

**Figure 3.**
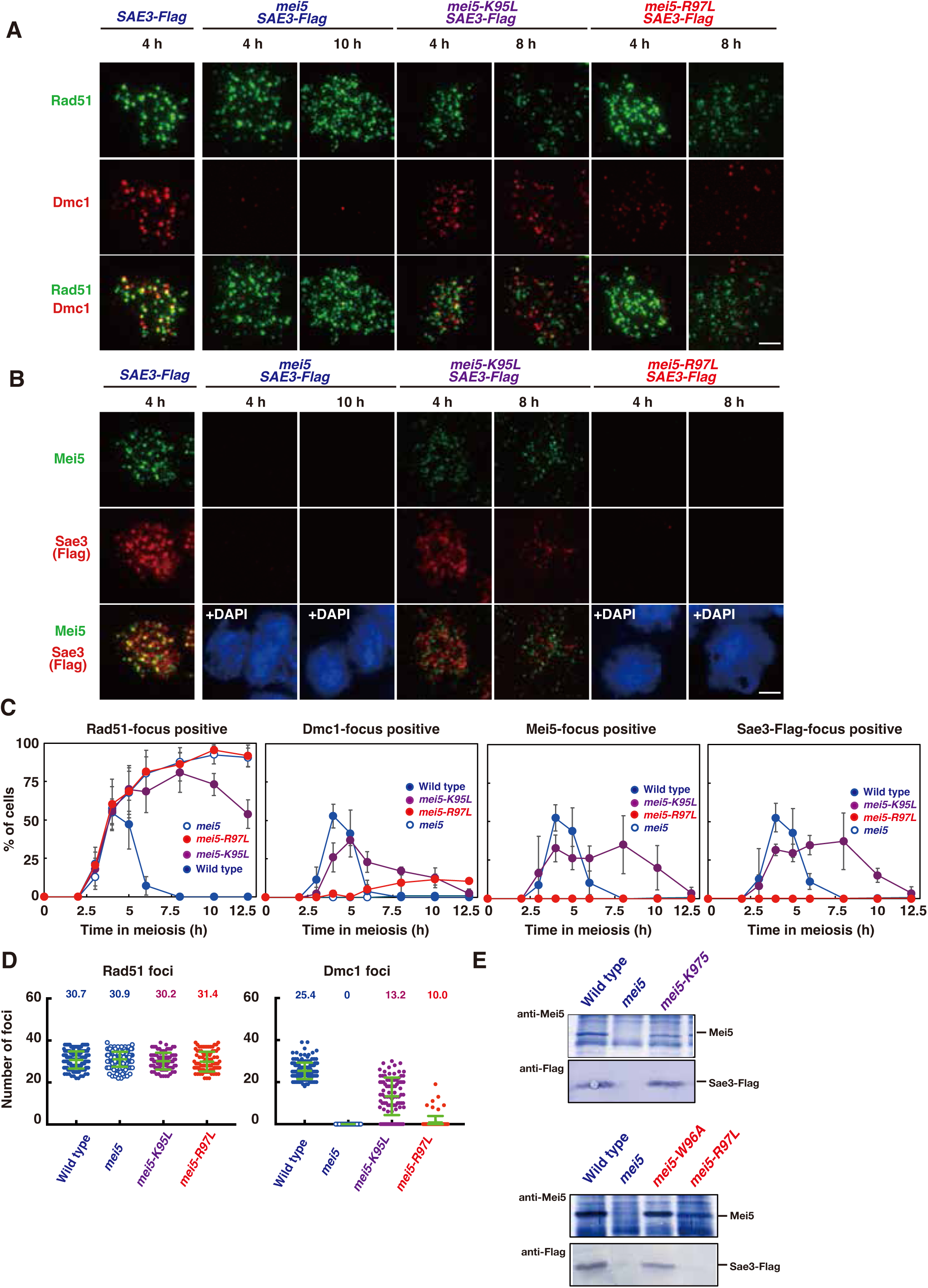
The *mei5-R97A* and *-R97L* are defective in Dmc1-assembly. A. Rad51 and Dmc1 staining. Nuclear spreads of *SAE3-Flag* cells with various *mei5* mutations were stained with anti-Rad51 (green), anti-Dmc1 (red), and DAPI (blue). Representative images at each time point under the two conditions are shown. Strains with *SAE3-FLAG* used are as follows: Wild-type, PSY31/32; *mei5::URA3* deletion, PSY166/167; *mei5-K95L*, PSY144/148; *mei5-R97L*, PSY157/158. Bar = 2 μm. B. Mei5 and Sae3 staining. Nuclear spreads were stained with anti-Rad51 (green), anti-Flag (Sae3, red), and DAPI (blue). Representative images at each time point under the two conditions are shown. Bar = 2 μm. C. Kinetics of assembly/disassembly of Rad51, Dmc1, Mei5 and Sae3. The number of cells positive for foci (with more than 5 foci) was counted at each time point. At each time point, more than 100 cells were counted. The average values and SDs of triplicates are shown. D. The number of foci of Rad51 and Dmc1 at 4 h was manually counted. The graphs show the focus number combined from three independent time courses. On the top, an average focus number in positive nucleus is shown. Error bars (green) is a mean with standard deviation. E. IP of Sae3-Flag by anti-Mei5 serum. Strains used are as follows: *SAE3-Flag*, PSY31/32; *mei5-K95L SAE3-Flag*, PSY144/148; *mei5-R97L SAE3-Flag*, PSY157/158.

We performed the immuno-precipitation (IP) of the Sae3-Flag protein in meiotic yeast cell lysates. From *SAE3-FLAG* cells, the IP using anti-Mei5 pulled down the Sae3-Flag protein efficiently (Figure 3E). On the other hand, the IP of Mei5-R97L did not reveal the Sae3-Flag protein while Mei5-W96A, which is the amino acid substitution of Trp96 next to Arg97, showed the complex formation with Sae3-Flag (Figure 3E). These showed that conserved Arg97 of Mei5 is critical for Mei5-Sae3 complex formation.

Although the *mei5-K95L* mutant behaves almost like wild-type (Figure 2), when combined with the *SAE3-Flag*, it showed a clear defect in the assembly and disassembly of Dmc1 (Figure 3A, C and Supplementary Figure 3A). The focus formation of Mei5 and Sae3-Flag is clearly reduced in the strain compared to the *SAE3-Flag* cells (Figure 3B, C, D, and Supplementary Figure S4). Mei5-K95L mutant protein can form a complex with Sae3-Flag (Figure 3E). These supports the idea that the C-terminal tagging of Sae3 affects the function and also that the *mei5-K95L* substitution weaken the function of Mei5(-Sae3).

### The *mei5-R117A* mutant produced a shorter Mei5 protein

Like the *mei5* null mutant, the *mei5-R117A* mutant, which is arrested in meiosis I (Figure 1B), is defective in Dmc1-focus formation but proficient in focus formation of Rad51 (Figure 4A, C). Rad51 foci persisted as meiosis progresses (Figure 4C). Immuno-staining of Mei5 revealed very few detectable Mei5 foci in *mei5-R117A* cells (Figure 4B, C). Arg117 is essential for the Mei5 function.

**Figure 4.**
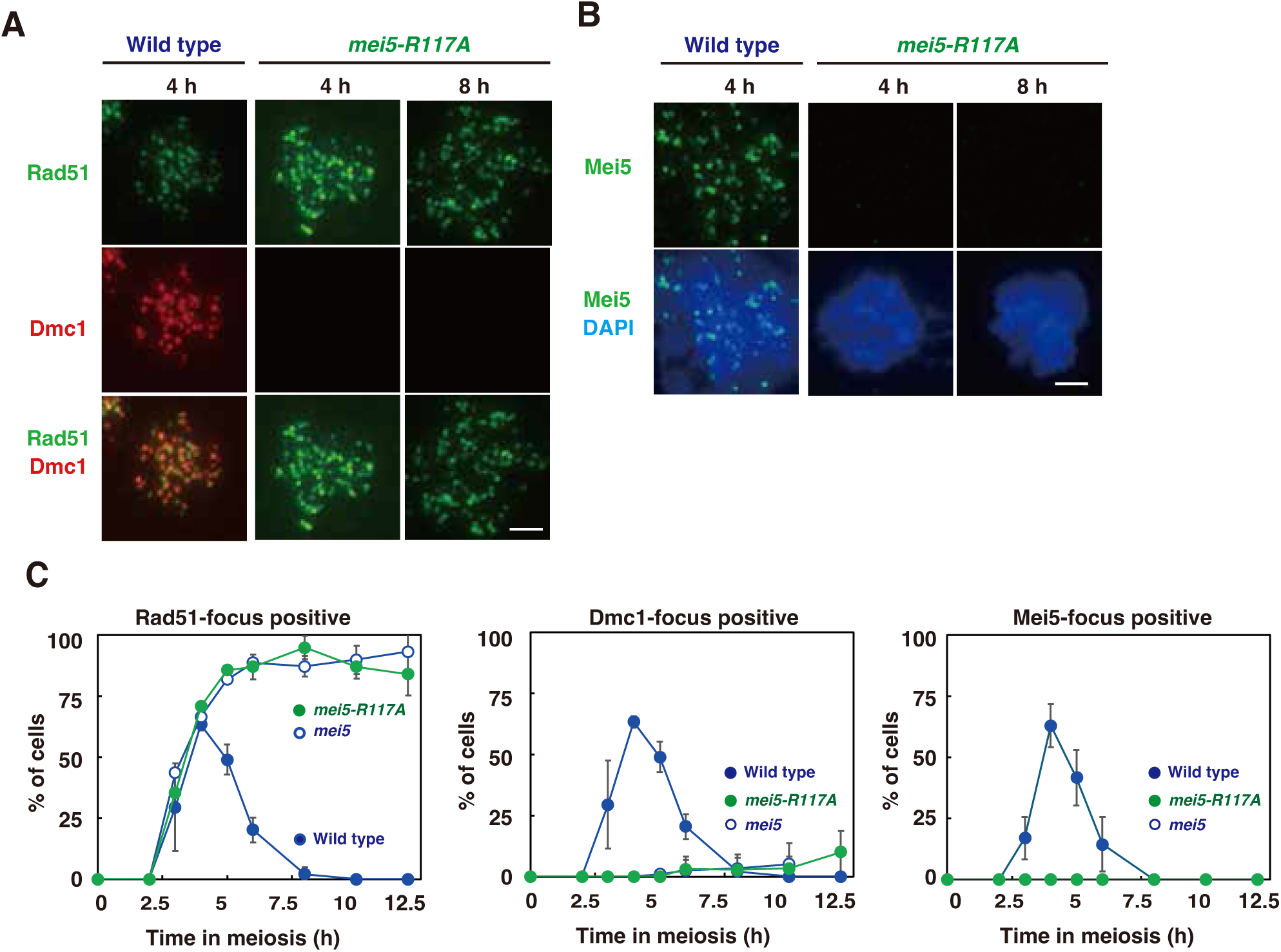
The *mei5-R117A* is defective in Dmc1 assembly. A. Rad51/Dmc1 staining in WT and *mei5-R117A* mutant cells. Nuclear spreads were stained with anti-Rad51 (green), anti-Dmc1 (red), and DAPI (blue). Representative images at each time point under the two conditions are shown. Strains used are as follows: Wild-type, NKY1551; *mei5-R117A*, SMY192/195. Bar = 2 μm. B. Mei5 staining. Nuclear spreads were stained with anti-Rad51 (red), anti-Mei5 (green), and DAPI (blue). Representative images at each time point under the two conditions are shown. Bar = 2 μm. C. Kinetics of Rad51/Dmc1/Mei5 foci in WT and *mei5-R117A* mutant cells. The number of cells positive for foci (with more than 5 foci) was counted at each time point. At each time point, more than 100 cells were counted. The average values and SDs of triplicates are shown.

On the other hand, on western blots using anti-Mei5 anti-serum, Mei5-R117A protein migrated as much faster bands at the position of around 25kD while wild-type protein (221 aa protein with calculated MW of 26 kD) ran at ∼30kD position (Figure 1C and 5A). Time course analysis showed that short-forms of Mei5-R117 protein as two bands appear at 2 h after the induction of meiosis, which is similar timing of the expression of the wild-type Mei5 protein. The shorter forms of Mei5-R117 protein are stable, which were detectable at late time points such as 10 h.

There are two possibilities for the formation of the shorter Mei5-R117 protein on the blot. First, wild-type Mei5 protein is post-translationally modified to the protein with a larger molecular weight than the unmodified protein. Indeed, Mei5 is SUMOylated at Lys56 (Bhagwat et al., 2021). Second, the mutant protein is subject to the processing after synthesized as a full-length protein. We compared the mobility of Mei5 protein expressed in meiotic yeast cells with purified Mei5(-Sae3) protein produced in *E. coli* and found the mobility of yeast Mei5 is similar to that Mei5 from *E. coli* (Figure 5A and C), suggesting that wild-type Mei5 protein is unlikely to be subject to the modification which induces the change of molecular weight on the gel. To confirm the processing of Mei5-R117A protein, we added the 3XFlag tag at the C-terminus of Mei5 protein. Wild-type Mei5-Flag protein was detected on blots as a band with a similar molecular weight (∼35kD) by using both anti-Mei5 and anti-Flag antibodies (Figure 5B). On the other hand, anti-Mei5 detected Mei5-Flag protein in *mei5-R117A-Flag* cell lysates, whose size is ∼20kD, similar to Mei5-R117A, indicating that the C-terminal Flag did not affect the size of Mei5-R117A protein. Moreover, the anti-Flag antibody could not the Mei5-R117A mutant protein with ∼20kD. These suggest that the C-terminal region is missing on the Mei5-R117A band. Moreover, there is a weak signal of full-length Mei5-R117A-Flag protein in the lysates, which is prominent at 6 h, implying the post-translational processing of a full-length Mei5-R117A-Flag protein. Of note, we often detected smaller bands of wild-type Mei5, which migrated almost similar two bands Mei5-R117A protein (red arrowhead in Figure 5A and B; the ratio of the two band are different between wild-type Mei5 and Mei5-R117A).

**Figure 5.**
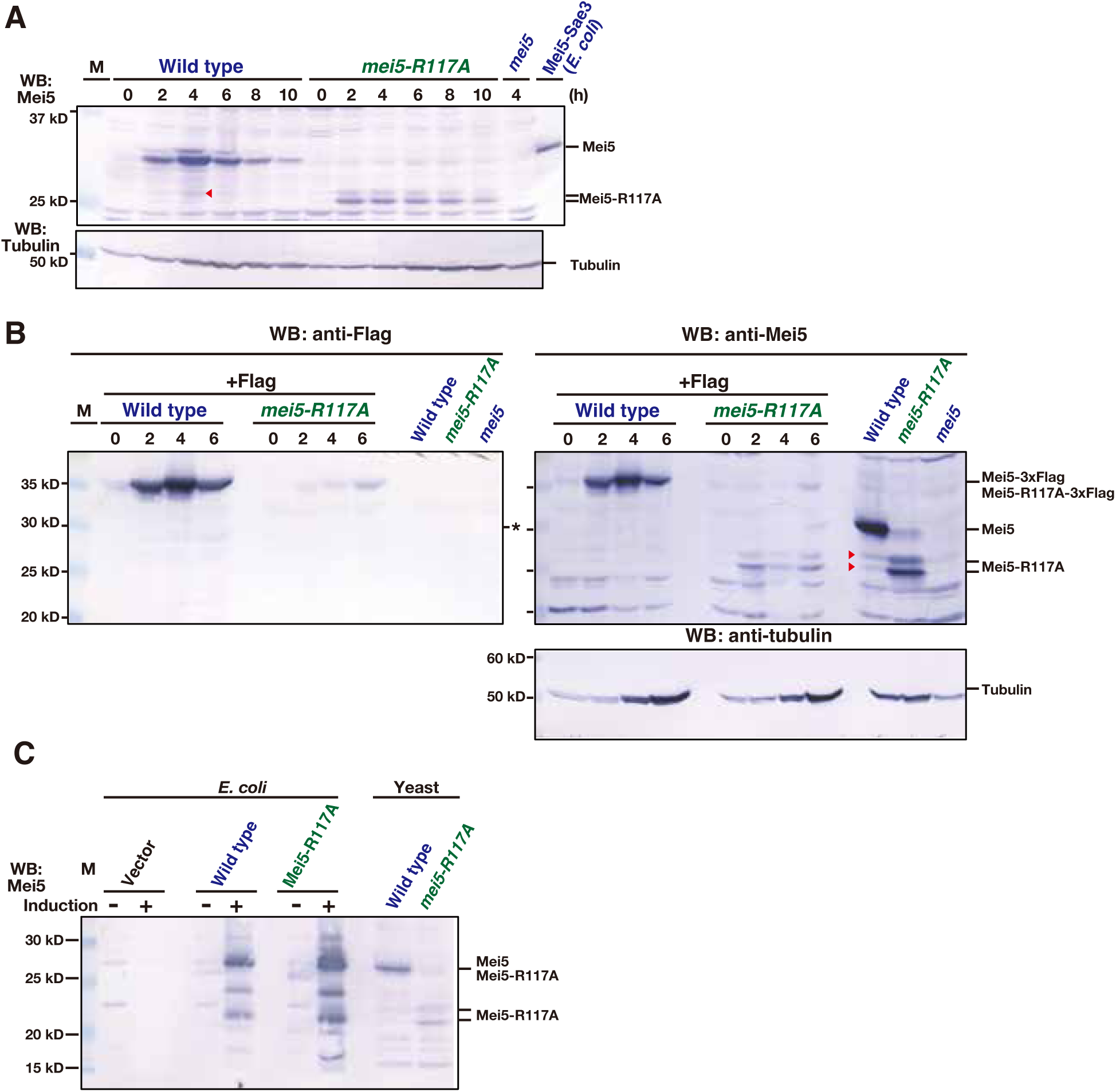
Mei5-R117A produces a truncated protein. A. Expression of Mei5-R117A. Cell lysates at each time were verified by western blotting with anti-Mei5 and anti-tubulin (control). Representative blots in duplicate are shown. Strains used are as follows: Wild-type, NKY1551; *mei5-R117A*, SMY192/195. A red arrowhead indicates a possible processed form of wild-type Mei5 protein. B. Expression of Mei5-R117A-3xFlag protein. Lysates of cell with Mei5-Flag or Mei5-R117A-Flag at each time were verified by western blotting with anti-Flag (left panel) anti-Mei5 (right panel), and anti-tubulin (bottom right). As a control, cells without the Flag tag on Mei5 were analyzed (right three lanes). Representative blots in duplicate are shown. Strains used are as follows: Wild-type with Mei5-Flag, SMY209/212; *mei5-R117A-Flag*, SMY210/222; Wild-type, NKY1551; *mei5-R117A*, SMY192/195. An asterisk means non-specific band cross-reacted with anti-Flag, which appear in late prophase I. Red arrowheads indicate a possible processed form of wild-type Mei5 protein. C. Expression of Mei5-R117A *in E. coli*. Cell lysates with or without the protein induction (with IPTG) were verified by western blotting with anti-Mei5. A representative blot in duplicate is shown. Vector, pET21a; wild-type, pET21a-Mei5-Sae3; pET21a-Mei5(R117A)-Sae3.

We expressed Mei5-R117A protein with Sae3 from *E. coli* cells and found the size of Mei5-R117A in *E. coli* lysates is similar to that of wild-type Mei5 protein (Figure 5A, C and Supplementary Figure S5). It is unlikely that the auto-cleavage activity of Mei5-R117A, supporting the idea that Mei5-R117A is subject to the processing in yeast.

### Mei5-R117A protein is sensitive to meiosis-specific posttranslational processing

To estimate the possible truncated position in the C-terminus of Mei5-R117A, we constructed the *mei5-d(190-221)* and *mei5-d(197-221)* mutants which lacked the C-terminal 32 and 25 aa, respectably, and compared the migration of these C-terminal truncated proteins with Mei5-R117A. Mei5-R117A migrated a bit faster than these truncated Mei5 proteins (Figure 6A and B). This suggests that the possible cleavage might occur around 190 residues of Mei5 (see blue region of a putative Mei5 structure in Figure 1A). Like the *mei5* deletion mutant, the *mei5-d(190-221)* and *mei5-d(197-221)* mutants arrest during meiotic prophase I (Figure 6C), indicating that the C-terminal two small α-helixes (α5 and α6) are critical for the Mei5 function.

**Figure 6.**
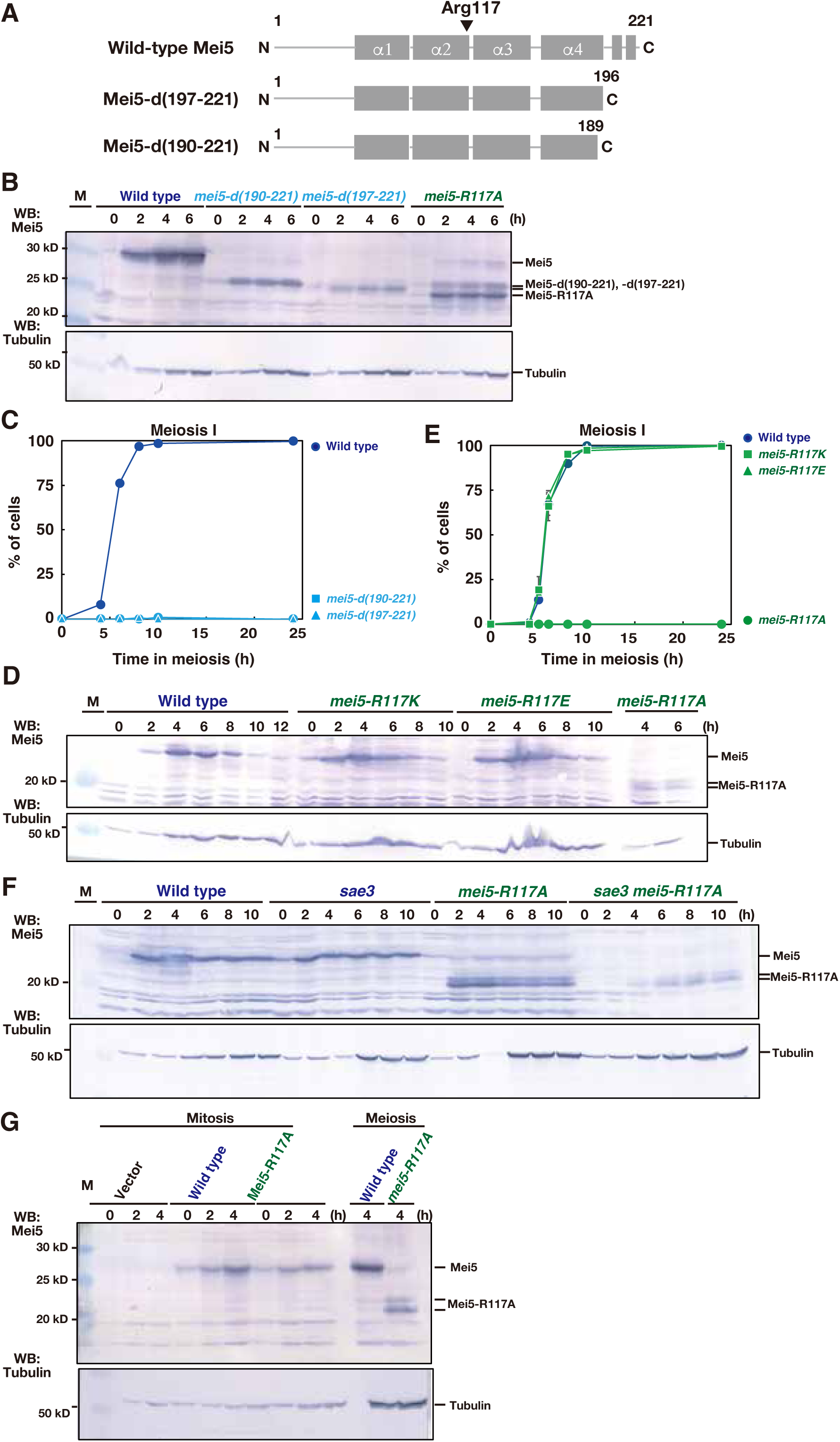
Mei5-R117A is processed post-translationally. A. Schematic drawing of Mei5 C-terminal deletion mutants; *mei5-d(190-221)* and *mei5-d(197-221)*. Rectangles present putative α-helixes. B. Expression of Mei5 C-terminal deletion mutants. The schematic drawing of the mei5 C-terminal deletion mutants (top). Gray rectangles represent α-helixes. Yeast cell lysates at each time were verified by western blotting (bottom) with anti-Mei5, and anti-tubulin (control). Representative blots in duplicate are shown. Strains used are as follows: Wild-type, NKY1551; *mei5-R117A*, SMY192/195; *mei5-d(190-221)*, SMY343/345; *mei5-d(197-221)*, SMY270/272. C. The entry into meiosis I in various strains was analyzed by DAPI staining. The number of DAPI bodies in a cell was counted. A cell with 2, 3, and 4 DAPI bodies was defined as a cell that passed through meiosis I. The graph shows the percentages of cells that completed MI or MII at the indicated time points. Wild-type, NKY1551; *mei5-R117A*, SMY192/195; *mei5-R117K*, SMY249/251; *mei5-R117E*, SMY253/255. D. Expression of the various Mei5-R117 mutant proteins. Representative blots in duplicate are shown. Strains used are as follows: Wild-type, NKY1551; *mei5-R117A*, SMY192/195; *mei5-R117K*, SMY249/251; *mei5-R117E*, SMY253/255. E. The entry into meiosis I in various strains was analyzed by DAPI staining. The average values and SDs of triplicates are shown. Wild-type, NKY1551; *mei5-R117A*, SMY192/195; *mei5-d(190-221)*, SMY343/345; *mei5-d(197-221)*, SMY270/272. F. Expression of Mei5-R117A in the *sae3* deletion. Lysates of cells with or without the *SAE3* were verified at each time by western blotting with anti-Mei5 and anti-tubulin (control). Representative blots in duplicate are shown. Strains used are as follows: Wild-type, NKY1551; *mei5-R117A*, SMY192/195; *sae3*, SMY261/263; *sae3 mei5-R117A*, SMY265/267. G. Expression of Mei5-R117A in mitotic yeast cells. The expression of Mei5 or Mei5-R117A protein from Gal1/10 promoter was verified by western blotting. At 0 h, 2% Galactose was added. Note-Gal1/10 did not work efficiently in SK1 background. Diploid cells with the expression vector of wild-type Mei5 (SMY291/292) and Mei5-R117A (SMY296/297) proteins were used. As a control, meiotic cell lysates of wild-type and *mei5-R117A* mutant cells at 4 h after meiosis induction were analyzed. Representative blots in duplicate are shown.

To know the requirement of the post-translational processing of Mei5-R117A, we constructed additional substitutions of Arg117 with lysine and glutamate, *MEI5-R117K* and *-R117E*, respectively. When we checked the size of these versions of Mei5, both Mei5-R117K and -R117E exhibited a similar size of the wild-type protein (Figure 6D). Indeed, in contrast to *mei5-R117A, MEI5-R117K* and *-R117E* cells showed normal MI progression with wild-type spore viability: wild-type, 97.1% (52 tetrads), *MEI5-R117K*, 97.5% (50 tetrads) and *MEI5-R117E*, 95.0% (5 tetrads: Figure 6E). Although we have not tested other substitutions of Arg117, the substitution to alanine is critical for the processing.

Given some protein processing in yeast depends on the proteasome, we treated the meiotic wild-type cells with a proteasome inhibitor, MG132 and the treatment with MG132 did not affect the mobility of Mei5-R117A as well as wild-type Mei5 (Supplementary Figure S6). The processing of Mei5-R117A seems to be independent of the proteasome.

We checked the complex formation of Mei5-R117A with Sae3 is necessary for the processing (Figure 6F). Like the *mei5-R117A* mutant, the *mei5-R117A* mutant with the *SAE3* deletion generated the truncated Mei5 protein. In the absence of Sae3, the Mei5-R117A protein is less stable relative to that in the presence of wild-type Mei5. Moreover, we expressed Mei5-R117A protein in mitotic yeast cells, which do not express Sae3, by putting it under the control of the *GAL1-10* promoter (which is defective for the induction in the SK1 strain). Mitotically-expressed Mei5-R117A protein migrated at the same position as wild-type Mei5 (Figure 6G). This suggests that the processing of Mei5-R117A protein is specific to meiosis.

## Discussion

Mei5-Sae3 promote Dmc1 assembly in the budding yeast. Previous biochemical studies show that Mei5-Sae3 promotes Dmc1-mediated D-loop formation by overcoming the inhibitory effect of RPA on Dmc1 assembly on the ssDNA (Chan et al., 2019; Cloud et al., 2012). How Mei5-Sae3 helps Dmc1 assembly on ssDNA remains largely unknown. The biochemical analysis of fission yeast and mouse Sfr1-Swi5, an ortholog of Mei5-Sae3, provides some mechanistic insight on how the complex promotes Rad51 assembly. Fission yeast and mouse Sfr1-Swi5 stabilizes ATP-bound form of Rad51 on ssDNAs (Haruta et al., 2006; Su et al., 2014). The structural analysis suggests that the fission yeast Sfr1-Swi5 bind to a groove of Rad51 filament (Kuwabara et al., 2012). Although we analyzed the role of conserved resides of Sae3 in the yeast meiosis (Sawant et al., 2023), the contribution of conserved amino acids in Mei5 has not been analyzed. Here, we characterized the substitution of basic amino acids on putative conserved α-helixes of Mei5 protein and identified key residues for *in vivo* Mei5 function.

### The role of conserved KWR motif in Mei5/Sfr1 family

The Mei5/Sfr1 orthologs are poorly conserved at the amino acid sequence. Among them, the KWK/R (Lys95-Trp96-Arg97) in the putative α2 helix in Mei5 are conserved among the family. In fission yeast Sfr1(-Swi5), the three residues are not involved in the binding with Swi5, rather are located on the surface of the complex (Supplementary Figure S1A). While the substitution of Lys85 did not affect Mei5 function *in vivo*, the substitution of Arg97 impaired the function. Although Mei5-R97L cannot form a complex efficiently *in vivo*, the protein might have a reduced Dmc1 assembly activity, suggesting a stable complex between Mei5 and Sae3 is not necessary for the function. Moreover, while Mei5-R97L (and -R97A) can recruit Dmc1 on meiotic chromosomes, Dmc1 assembly proceeds Mei5 assembly in the *mei5-K95A/L* mutant. This suggests that a stoichiometric amount of Mei5-Sae3 relative to Dmc1 is not necessary for the role of the complex as the Dmc1 mediator. In the other word, a small amount of Mei5-Sae3 is sufficient for Dmc1 assembly.

The *mei5-K95L* mutant alone is almost normal in Dmc1 assembly. On the other hand, the combination of the substitution with the *SAE3-Flag* makes this Mei5-Sae3 complex defective in the assembly of Dmc1 as well as the loading of Mei5 and Sae3 (Figure 3C). This suggests a possible role of Arg85 in Mei5 function. The position of KWR motif is near the C-terminus of fission yeast Swi5 on the predicted structure (Supplementary Figure S1).

In wild type, the loading of Dmc1 and Mei5-Sae3 is mutually inter-dependent (Hayase et al., 2004; Tsubouchi & Roeder, 2004). Interestingly, Mei5-R97A/L binds on meiotic chromosomes later than Dmc1, suggesting that the loading of Mei5-Sae3 is temporally separable from Dm1 loading. Moreover, later Mei5-Sae3 seems to be able to bind Dmc1 ensembles. This implies the role of Mei5-Sae3 in post-assembly stage of Dmc1 filament such as the stabilization of the filament.

### Possible post-translational processing of Mei5-R117A protein

Arg117 substitution with Ala but not Lys or Glu induced the post-translational cleavage of the mutant Mei5 protein, which could generate Mei5 protein deleted for the C-terminal ∼30 amino acids. Since the two C-terminal deletion *mei5* mutants, *mei5-d(190-221)* and *mei5-d(197-221)*, are defective in meiosis, we speculate this processing inactivates the Mei5 function. Although we do see the robust cleavage of Mei5-R117A protein *in vivo*, we do not have clear evidence that wild-type Mei5 is a target for the similar post-translational processing. In wild-type cells, we detected that Mei5 protein with a similar size to processed Mei5-R117A in addition to full-length Mei5, although the amount of the processed proteins is relatively to very small compared to full-length protein. This suggest that the wild-type Mei5 is subject to the similar processing to Mei5-R117A protein, although less sensitive compared to the mutant protein So even if such the processing may exist, it does not seem to operate under normal condition of meiosis.

The processing of Mei5-R117A is rapid and under the control of biologically regulated process, since we see the cleavage of the mutant Mei5 only in meiotic cells, but not in mitotic cells. There are limited reports on post-translational processing of the protein in the budding yeast, although such the phenomenon is often reported in multi-cellular organism. In the budding yeast, the membrane-bound Mga2 transcriptional factor p120 is converted into un-anchored Mga2p90, which is imported to the nucleus for the transcriptional regulation in mitotic cells (Bhattacharya et al., 2009; Hoppe et al., 2000). This processing depends on the proteasome and Cdc48 unfoldase. In meiosis, nuclear envelope protein, Mps3, is cleaved in its N-terminal ∼90 amino acids by the proteasome-dependent mechanism only in a specific window of meiotic prophase I (Li et al., 2017). The cleavage of Mei5-R117A might be related to these reported processes. However, the proteasome inhibitor, MG132, treatment did not inhibit the appearance of Mei5-R117A. It is likely that the processing of Mei5-R117A is proteasome-independent. To elucidate the biological role of this processing as well as to get the mechanical insight, the identification of Mei5 residues (or regions) necessary for the cleavage or the cleavage site(s) as well as the protease mediating the process is necessary.

### Experimental Procedures Strains

All strains described here are derivatives of SK1 diploids, NKY1551 (*MATα/MAT**a**, lys2/”, ura3/”, leu2::hisG/”, his4X-LEU2-URA3/his4B-LEU2, arg4-nsp/arg4-bgl*). The genotypes of each strain used in this study are described in Supplemental Table S1.

### Strain Construction

The *mei5* mutant genes were constructed by the pop-in/pop-out methods. PCR-based site-directed mutagenesis was carried out using the yIPlac195 plasmid with wild-type *MEI5* gene as a template. The sequence of primer DNAs using site-directed mutagenesis is provided in Table S2. DNA changes were confirmed by DNA sequencing. The plasmids with the mutant *MEI5* gene were introduced in the yeast strain of MSY832 by transformation with Li acetate and yeast cells were selected for uracil prototroph. The presence of the mutants was confirmed by the restriction digestion of PCR products. The candidate yeast cells were grown overnight in YPAD liquid culture and were plated on a selection media containing 5-FOA (5-fluoroortic acid) for uracil auxotroph. The C-terminal *mei5* deletion mutants were constructed by PCR-mediated one-step replacement using pFA6a-KamMX4. The mutations were confirmed again by restriction digestion of PCR products and the sequencing.

### Anti-serum and antibodies

Mouse anti-Flag (anti-DYKDDDDK tag; Wako 012-22384), rabbit anti-Mei5 serum (Hayase et al., 2004), and anti-tubulin (MCA77G, Bio-Rad/Serotec, Ltd) were used for western blotting. Guinea pig anti-Rad51 (M. Shinohara, Gasior, Bishop, & Shinohara, 2000), rabbit anti-Rfa2 (Hayase et al., 2004), rabbit anti-Dmc1 (Sasanuma et al., 2013), and rabbit anti-Mei5 serum (Hayase et al., 2004) were used for staining. With a higher background, IgG was purified using the IgG purification kit (APK-10A, Cosmo Bio co. Ltd). The secondary antibodies for staining were Alexa-488 (Goat) and -594 (Goat) IgG used at a 1/2000 dilution (Themo Fishers).

### Meiotic Time course

*Saccharomyces cerevisiae* cells were patched onto YPG plates (2% bacteriological peptone, 1% yeast extract, 3% glycerol, 2% agar) and incubated at 30 °C for 12 h. Cells were spread into YPD plates (2% bacteriological peptone, 1% yeast extract, 2% glucose, 2% agar) and grown for 48 h to isolate a single colony. The single colony was inoculated into 2 ml of YPD liquid medium and grown to saturation at 30 °C overnight. To synchronize cell cultures, the overnight cultures were transferred to a SPS medium (1% potassium acetate, 1% bacteriological peptone, 0.5% yeast extract, 0.17% yeast nitrogen base with ammonium sulfate and without amino acids, 0.5% ammonium sulfate, 0.05 M potassium biphthalate) and cells were grown for 16–17 h. Meiosis was induced by transferring the SPS-cultured cells to a pre-warmed SPM medium (1% potassium acetate, 0.02% raffinose). The cells were collected at various times after being transferred to the SPM medium.

### Immuno-staining

Immunostaining of chromosome spreads was carried out as described previously (M. Shinohara et al., 2000; M. Shinohara, Sakai, Ogawa, & Shinohara, 2003). Briefly, spheroplasts burst in the presence of 1% paraformaldehyde and 0.1% lipsol. Stained samples were observed using an epi-fluorescent microscope (BX51; Olympus/Evident) with a 100 X objective (NA1.3). Images were captured by CCD camera (Cool Snap; Roper) and, then processed using iVision (Sillicon), and Photoshop (Adobe) software. For focus counting, ∼30 nuclei were counted at each time point.

### Western blotting

Western blotting was performed as described previously (Hayase et al., 2004; M. Shinohara, Oh, Hunter, & Shinohara, 2008). Western blotting was performed for cell lysates extracted by the TCA method. After being harvested and washed twice with 20% TCA, cells were roughly disrupted with zirconia beads by the Multi-beads shocker (Yasui Kikai Co Ltd, Japan). Protein precipitation recovered by centrifugation was suspended in SDS-PAGE sample buffer (pH 8.8) and then boiled at 95°C for 10 min. After the electrophoresis, the proteins were transferred onto the Nylon membrane (Immobilon-P, Millipore) and incubated with primary antibodies in a blocking buffer (1X PBS, 0.5% BSA) and then alkaline phosphatase-conjugated secondary antibody (Promega, US). The color reaction was developed with NBT/BCIP solution (Nacalai Tesque, Japan).

### Pulsed-field gel electrophoresis

For pulsed-field gel electrophoresis (PFGE), chromosomal DNA was prepared in agarose plugs as described in (Bani Ismail, Shinohara, & Shinohara, 2014) and run at 14 °C in a CHEF DR-III apparatus (BioRad) using the field 6V/cm at a 120°angle. Switching times followed a ramp from 15.1 to 25.1 seconds. The duration of electrophoresis was 41 h for all chromosomes. For short chromosomes, 5 to 30 seconds and others for 20 to 60 seconds.

### Immuno-precipitation

Yeast cells were resuspended in the Lysis buffer (50 mM HEPES-NaOH [pH 7.5], 140 mM NaCl, 10% Glycerol, 1 mM EDTA, 5% NP-40). An equal amount of glass beads (Zircona Y2B05) was added along with the Protease inhibitor cocktail 10X: Roche “Complete™, Mini, EDTA-free Protease Inhibitor Cocktail”; 4693159001. The cells were disrupted using the Multi-beads shocker (Yasui Kiki; 2300 rpm, 60 sec on; 60 sec off cycle, 4 times). The lysates were incubated with magnetic beads (Dynal M260 Protein-A conjugated; GE Healthcare) coated with the anti-Mei5 antibody (6 μl serum in 100 μl beads) at 4 °C for 12 h and washed extensively (Sasanuma et al., 2013). Bound proteins were eluted by adding the SDS sample buffer and were analyzed on an SDS-PAGE gel, transferred to a nylon membrane (Millipore Co. Ltd), and probed with specific antibodies.

### Protein expression in *E. coli*

The expression plasmid for Mei5-R117A-Sae3 was generated from pET21a-Mei5-Sae3 (Hayase et al., 2004) by PCR amplification and Gibson assembly method using primers (5’-CTTTAATCAAAATCAATGCAATGGGCGGCTATAAAGAT-3’ and 5’-ATCTTTATAGCCGCCCATTGCATTGATTTTGATTAAAG-3’). Resulting pET21a-Mei5-R117A-Sae3 plasmid was introduced into BL21(DE3) with pLysS and the protein expression was induced at 30°C for 3 h by the addition of IPTG (final 1 mM). The protein expression in whole cell lysate was analyzed by SDS-PAGE followed by CBB staining or Western blotting.

### Software and Statistics

Figures for protein structure analysis were generated by PyMOL. Means ± S.D values are shown. Datasets (focus number) were compared using the Mann-Whitney U-test (Prism, GraphPad).

## Acknowledgments

We are grateful to members of the Shinohara lab. S.M.W., O. P. O., and P.S. was supported by a Japanese government scholarship and by the Institute for Protein Research. This work was supported by Osaka Fermentation Foundation to A.S.

## Author contributions

A.S. conceived and designed the experiments. S.M.W. and P.S. carried out yeast experiments. O.P.O. constructed some strains. A.F. expressed proteins. S.M.W., P.S., Y.F., M.I., and A.S. analyzed the data. A.S. wrote the manuscript with inputs from others.

---

**Table S1.**
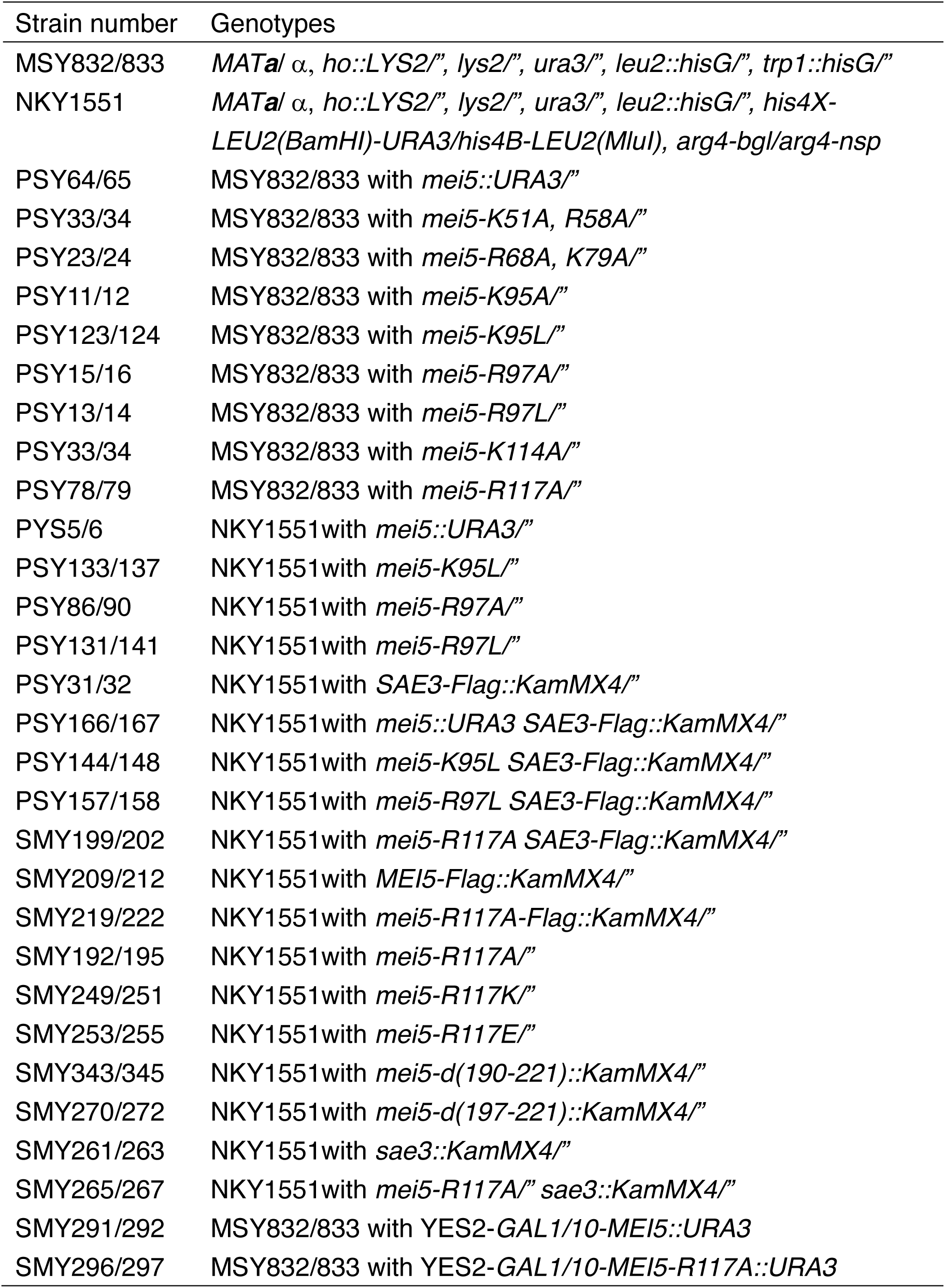
Strain List

**Table S2.**
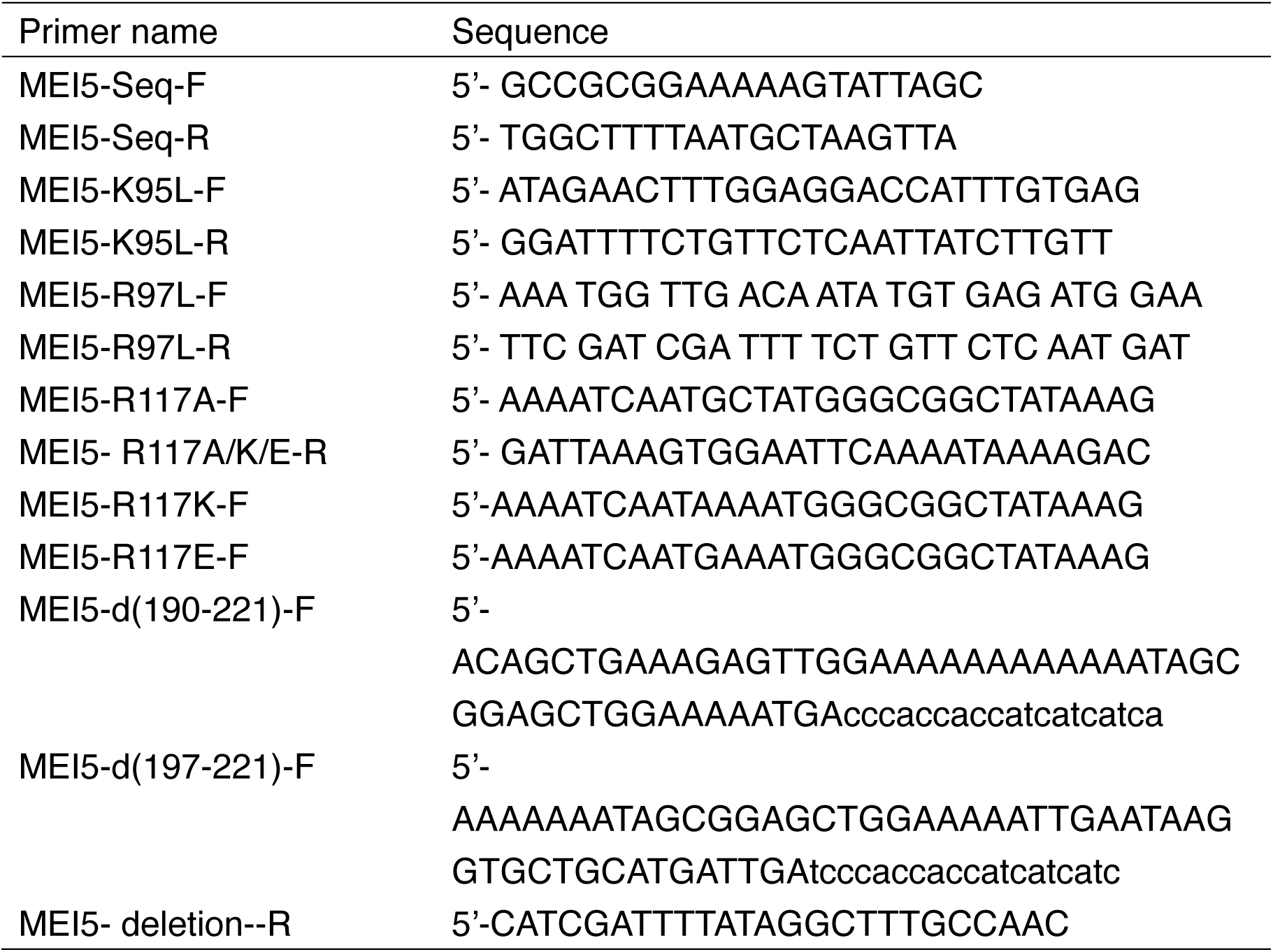
Primer List

**Supplemental Figure S1.**
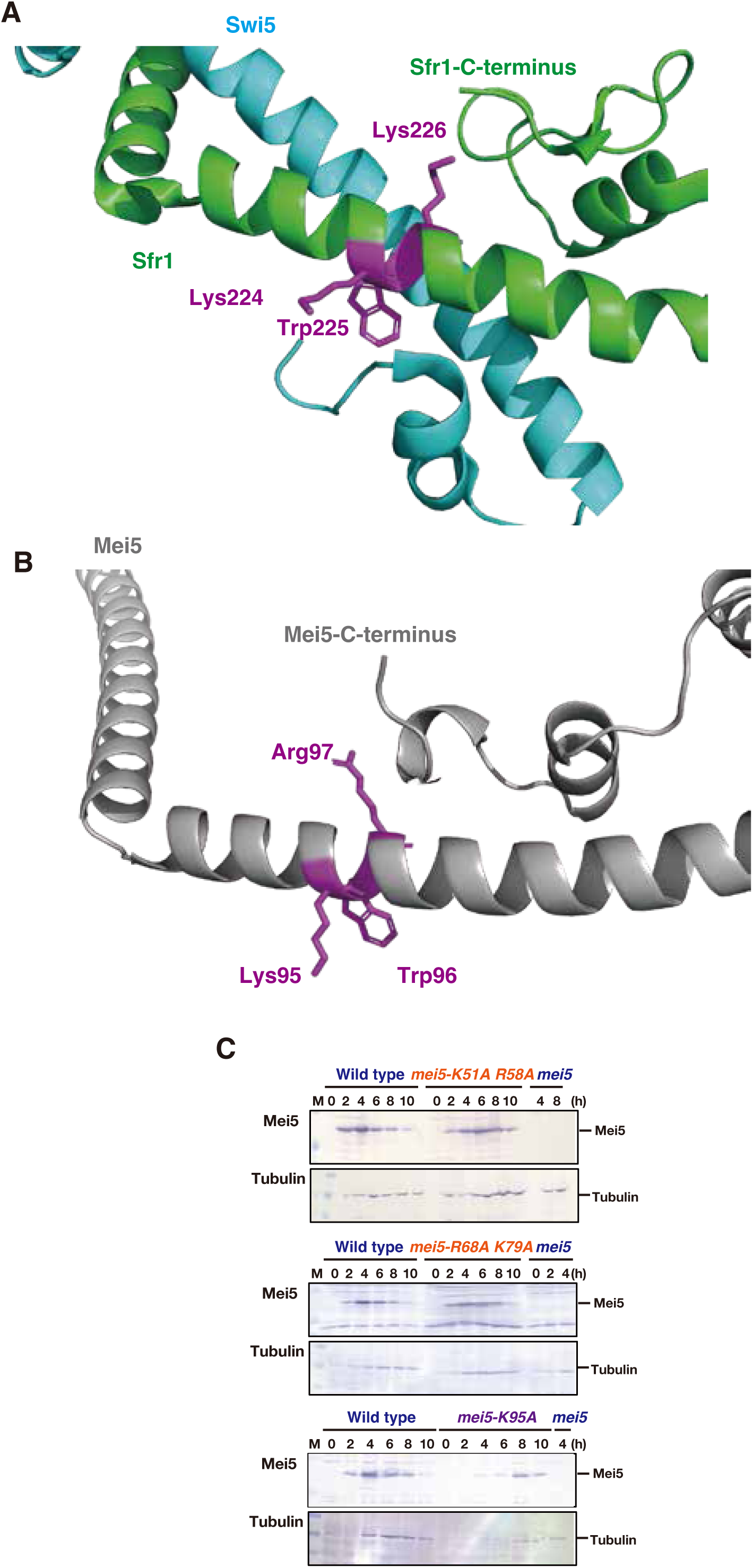
A. Crystal structure of fission yeast Sfr1-Swi5. An enlarged portion of Sfr1 (green) and Swi5 (pale blue) are shown. Conserved KWK/R residues are shown in the stick model (purple). B. AlphaFold2-predicted Mei5 structure. Conserved KWK/R residues; Lys95, Trp96, and Arg97, are shown in the stick model (purple). C. Expression of various mutant Mei5 proteins in meiosis. Lysates obtained from the cells at various time points during meiosis were analyzed by western blotting using anti-Mei5 (upper) or anti-tubulin (lower) antibodies.

**Supplemental Figure S2.**
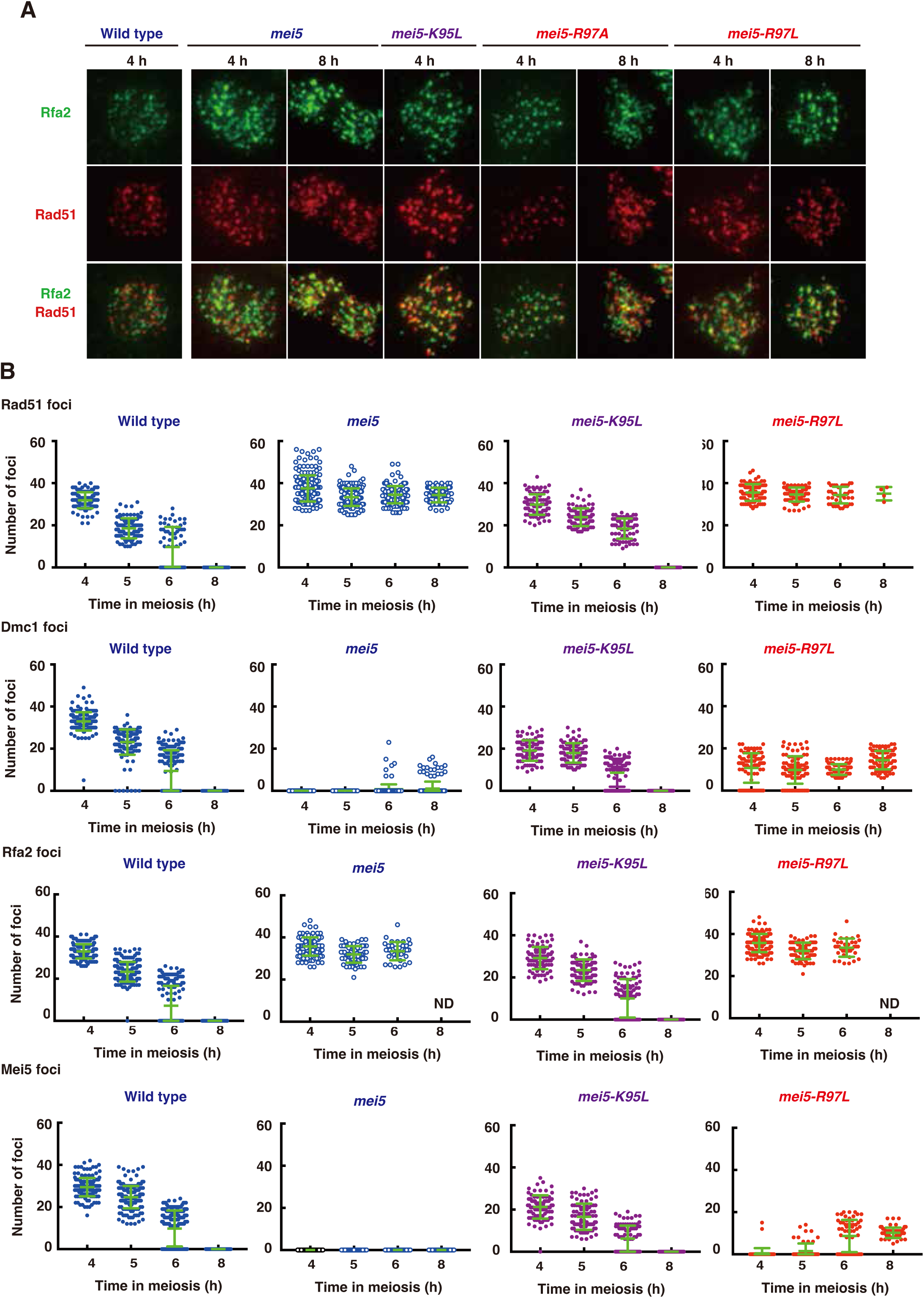
A. Rad51 and Rfa2 staining. Nuclear spreads were stained with anti-Rad51 (red), anti-Rfa2 (green), and DAPI (blue). Representative images at each time point under the two conditions are shown. Bar = 2 μm. B. Focus number at different time points: At each time point, foci were manually counted. The graphs show the focus number combined from three independent time courses. Error bars (green) is a mean with standard deviation.

**Supplemental Figure S3.**
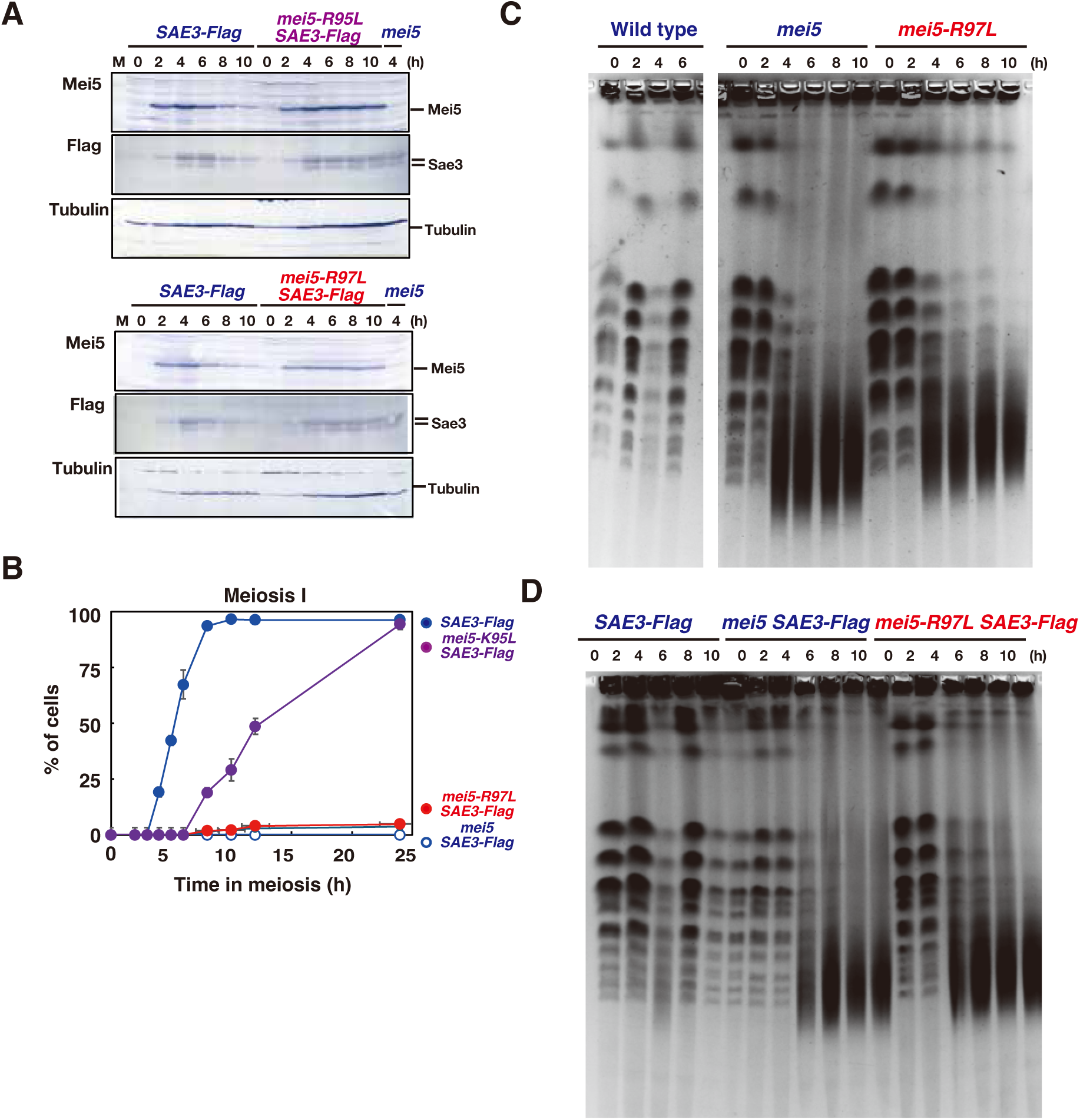
A. Expression of various mutant Mei5 and Sae3-Flag proteins in meiosis. Lysates obtained from the cells at various time points during meiosis were analyzed by western blotting using anti-Mei5 (upper), anti-Flag (middle) or anti-tubulin (lower) antibodies. B. The entry into meiosis I in various strains was analyzed by DAPI staining. The number of DAPI bodies in a cell was counted. A cell with 2, 3, and 4 DAPI bodies was defined as a cell that passed through meiosis I. The graph shows the percentages of cells that completed MI or MII at the indicated time points. C. CHEF analysis of meiotic DSB repair. Chromosomal DNAs from Wild-type (NKY1551), *mei5*(PSY5/6), and *mei5-R97L* (PSY131/141) cells were studied by CHEF electrophoresis. D. CHEF analysis of meiotic DSB repair. Chromosomal DNAs from *SAE3-Flag* (PSY31/32), *mei5 SAE3-Flag* (PSY166/167), and *mei5-R97L SAE3-Flag* (PSY157/158) cells were studied by CHEF electrophoresis.

**Supplemental Figure S4.**
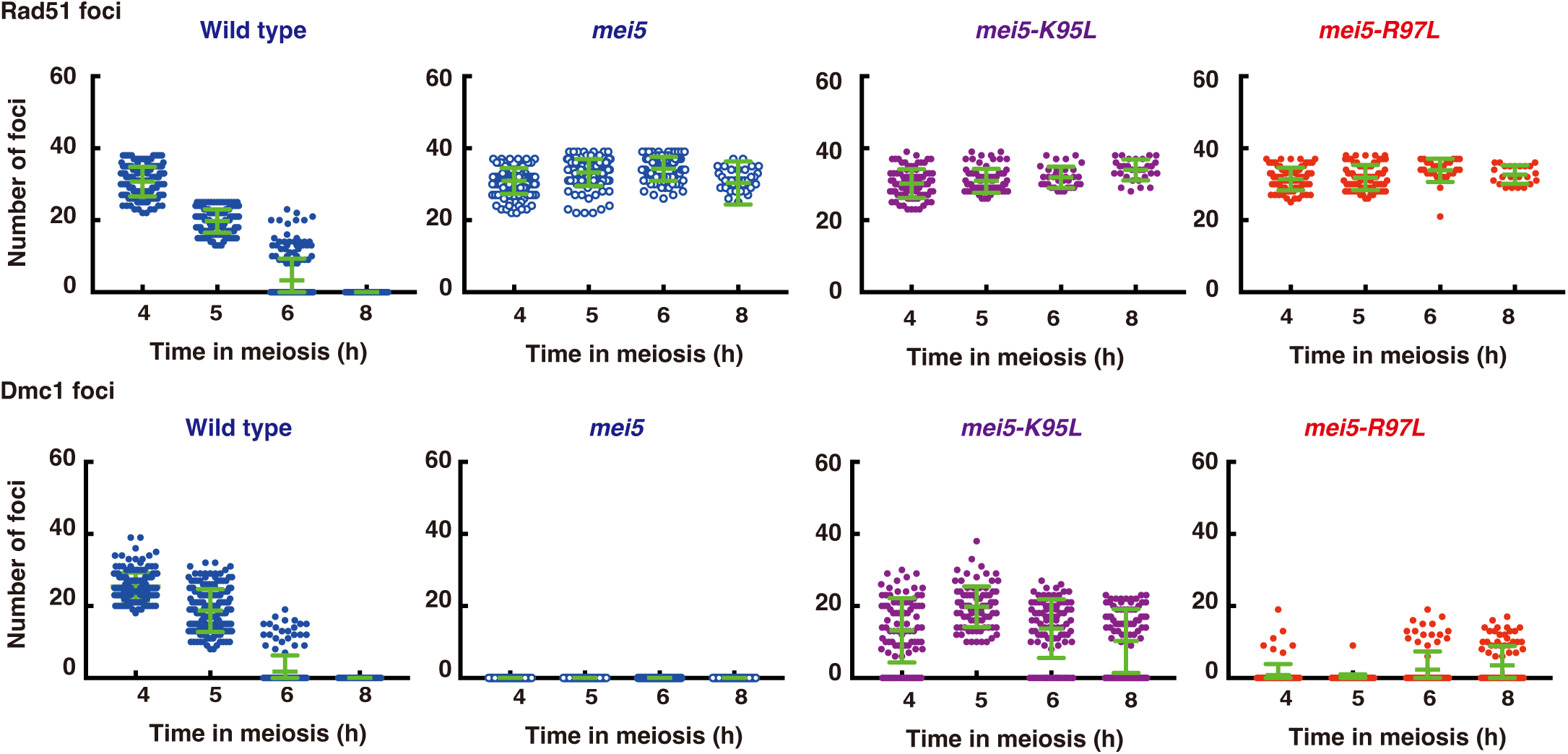
Focus number at different time points: At each time point, foci were manually counted. The graphs show the focus number combined from three independent time courses. Error bars (green) is a mean with standard deviation.

**Supplemental Figure S5.**
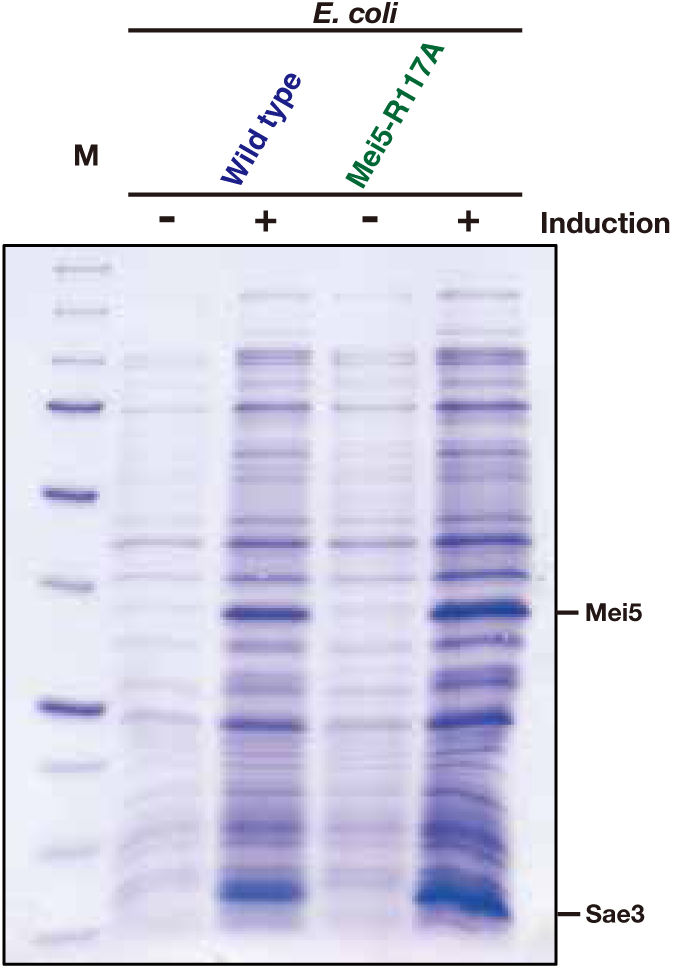
E. *coli* cell lysates were stained with Coomassie Brilliant blue. BL21(DE3) cells with various vectors were cultured with or without IPTG for 3 hours.

**Supplemental Figure S6.**
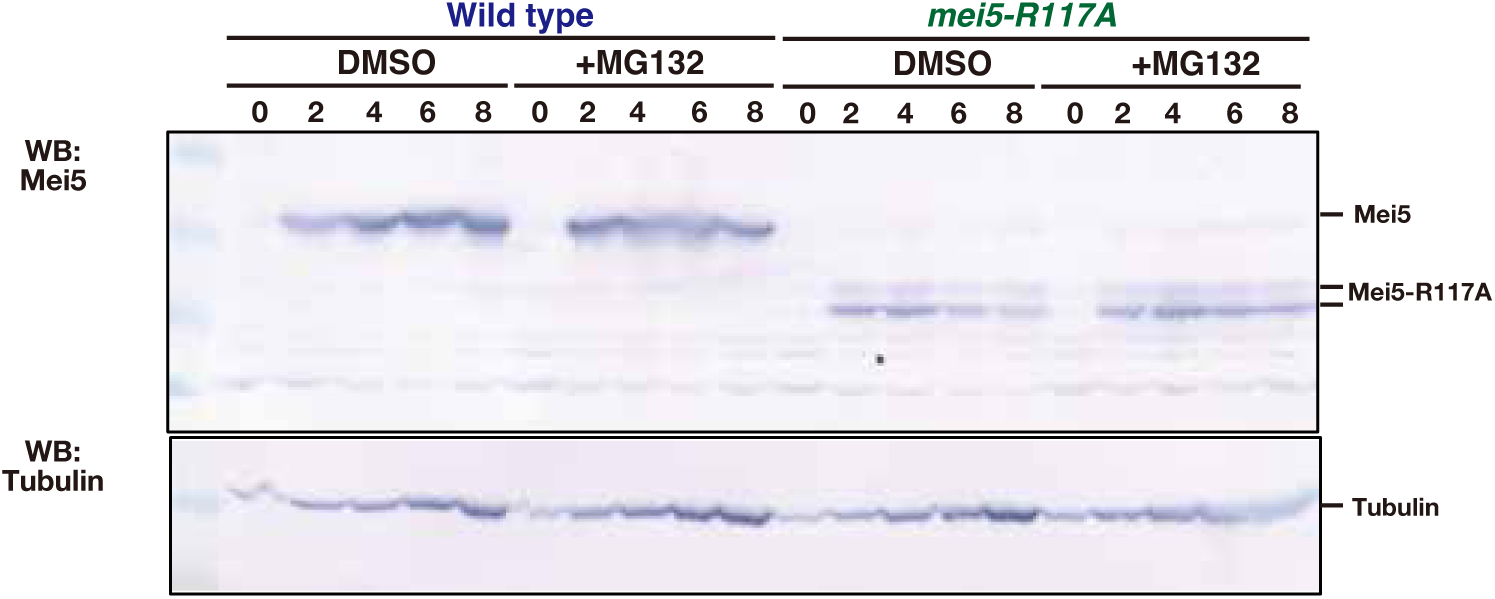
Expression of Mei5-R117A. Meiosis was induced in the presence or the absence of a proteasome inhibitor, MG132. Cell lysates at each time were verified by western blotting with anti-Mei5 and anti-tubulin (control). MG132 was added at 0 h at a concentration of 50 μM. Strains used are as follows: Wild-type, NKY1551; *mei5-R117A*, SMY192/195.

